# Sugar restriction and blood ingestion shape divergent immune defense trajectories in the mosquito *Aedes aegypti*

**DOI:** 10.1101/2023.01.24.525229

**Authors:** Dom Magistrado, Noha K. El-Dougdoug, Sarah M. Short

## Abstract

Immune defense is comprised of 1) resistance: the ability to reduce pathogen load, and 2) tolerance: the ability to limit the disease severity induced by a given pathogen load. The study of tolerance in the field of animal immunity is fairly nascent in comparison to resistance. Consequently, studies which examine immune defense comprehensively (i.e., considering both resistance and tolerance in conjunction) are uncommon, despite their exigency in achieving a thorough understanding of immune defense. Furthermore, understanding tolerance in arthropod disease vectors is uniquely relevant, as tolerance is essential to the cyclical transmission of pathogens by arthropods. Here, we tested the effect(s) of dietary sucrose concentration (high or low) and blood meal (present or absent) on resistance and tolerance to *Escherichia coli* infection in the yellow fever mosquito *Aedes aegypti*. Resistance and tolerance were measured concurrently and at multiple timepoints. We found that both blood and sucrose affected resistance. Mosquitoes from the low sugar treatment displayed enhanced resistance at all timepoints post-infection compared to those from the high sugar treatment. Additionally, blood-fed mosquitoes showed enhanced resistance compared to non-blood-fed mosquitoes, but only on day 1 post-infection. Sucrose had no effect on tolerance, but the effect of blood was significant and dynamic across time. Specifically, we show that consuming blood prior to infection ameliorates a temporal decline in tolerance that mosquitoes experience when provided with only sugar meals. Taken together, our findings indicate that different dietary components can have unique and sometimes temporally dynamic impacts on resistance and tolerance. Finally, our findings 1) highlight the value of experimental and analytical frameworks which consider the explicit testing of effects on both resistance and tolerance as separate, but equally important, components of immune defense, and 2) underscore the importance of including a temporal component in studies of immune defense.

## Introduction

An organism’s response to infection (i.e., immune defense) is comprised of both resistance and tolerance (1,2). Resistance is defined as the ability to reduce the number of pathogens inside the body, while tolerance is defined as the ability to limit the impact of the infection on host fitness. Both strategies are critical to surviving infection. Resistance and tolerance were first conceptualized by botanists in the late 1800s (3), and plant biologists have obtained major insights into both resistance and tolerance in years since. In contrast, scientists studying animal hosts have focused disproportionately on resistance until recently (2,4,5). As a result, immune tolerance in animals is a nascent, but quickly developing, field of study. Tolerance strategies likely include tactics such as repair of pathogen-induced tissue damage, detoxification of pathogen by-products, limitation of immune response-mediated self-injury (i.e. immunopathology), and general homeostasis promotion during and following infection – all strategies that promote the health of the host without necessarily affecting pathogen levels (2,6–9). Ecological immunology posits that resistance and tolerance, as components of host defense, are costly and should only be employed when the benefits outweigh the costs (10). Therefore, a host’s balance between resistance and tolerance is likely important from a resource limitation perspective, as an inappropriate balance between resistance and tolerance may lead to an undesirable infection outcome. For example, a strong resistance response may result in complete pathogen elimination, but such an outcome could leave inadequate energy reserves for repairing infection-induced damages. If this results in reduced lifetime reproductive fitness, it would not be a successful strategy.

While an appropriate equilibrium between resistance and tolerance is important in all host-pathogen systems, it has unique relevance in hematophagous arthropods that transmit pathogens. Arthropod-borne pathogens are highly diverse, and include viruses, bacteria, protozoan parasites, and filarial worms. Diseases caused by these pathogens comprise more than 17% of all global infectious diseases (11), and thus impose a massive public and veterinary health burden. Arthropod-borne pathogens are ingested by the arthropod when it takes a blood meal from an infected vertebrate host. Because transmission is categorically dependent upon the arthropod’s ability to tolerate infection long enough to spread the pathogen to a subsequent vertebrate host, studying immune tolerance is critical to the ability to understand and address arthropod-borne pathogen transmission. We therefore explored the effect of diet on resistance and tolerance to infection concurrently in the yellow fever mosquito, *Ae. aegypti*, which transmits multiple human pathogens including dengue virus and Zika virus.

Female adult mosquitoes have evolved to consume two meal types: 1) nectar, which is rich in sugar, and 2) blood, which is rich in protein and necessary for egg production in most species (12,13). Consumption, storage, and digestion differ appreciably between the two meal types. For example, sugar meals are stored in the ventral diverticulum (crop) while blood meals bypass the crop and are directly sent to the midgut. Sugar meals can be stored in the crop until needed for energy-intensive activities such as flight while blood digestion typically begins within a few hours of feeding (14). Blood digestion, unlike sucrose digestion, is also associated with significant physiological stress due to rapid shifts in temperature and pH, gut distension, heme toxicity resulting from hemoglobin digestion, and midgut redox stress (15–21).

Diet has previously been implicated in immune defense in arthropods. For example, lower dietary sugar concentrations have been shown to enhance resistance to bacterial infection in *Drosophila melanogaster* and resistance to *Plasmodium* infection in *Anopheles stephensi* (21,22). Effects of diet on tolerance have been explored as well. For example, genotype interacts with diet to impact tolerance to bacterial infection in *D. melanogaster* (22). Additionally, a study that investigated the effects of a blood meal on bacterial infection in *Ae. aegypti* found that blood impacts resistance and tolerance early in infection, but they also found that the effects were dose-dependent (23). In the present study, we tested the effects of two adult female *Ae. aegypti* diet components, 1) low/high sucrose diet concentration and 2) the ingestion of a blood meal, on resistance and tolerance to bacterial infection. We measured resistance and tolerance concurrently at multiple timepoints across a 5-day infection timecourse, allowing us to examine each treatment group’s resistance/tolerance trajectory, as well as test for any potential interactions between the two components of the mosquito’s diet in affecting immune defense. Our results contribute to the understanding of how arthropods of medical importance withstand pathogen infection, and more specifically, elucidate effects of blood feeding and sugar feeding on immune defense that may be conserved amongst other hematophagous arthropod vectors.

## Results

We investigated the effect of dietary sucrose concentration and blood feeding on resistance and tolerance to infection over time. To accomplish this, we exposed female *Ae. aegypti* mosquitoes to four different diet treatments: 10% sucrose alone, 10% sucrose + blood meal, 1% sucrose alone, and 1% sucrose + blood meal. We then infected them with *E. coli* (S17) and measured bacterial load and survival at multiple timepoints post-infection (Fig 1). The resultant dataset comprises seven variables (Table 1) and was used to build multiple models describing the effect(s) of both Blood and Sucrose on resistance and tolerance to bacterial infection across time.

**Table 1.** Description of terms included in models.

| *Variable* | *Type* | *Description* |
| --- | --- | --- |
| Survival (Alive, Dead) | Weighted proportion following a binomial distribution | Proportion of mosquitoes alive, weighted for sample size |
| Blood | Categorical – binary | Mosquitoes either received or did not receive a blood meal |
| Sucrose | Categorical – binary | Mosquitoes received either 1% sucrose or 10% sucrose |
| Day | Discrete | Day (1, 3, or 5) post-infection at which measurements were taken |
| Bacterial Load | Continuous | Median Colony Forming Units per mosquito at time of sampling |

**Fig 1.**
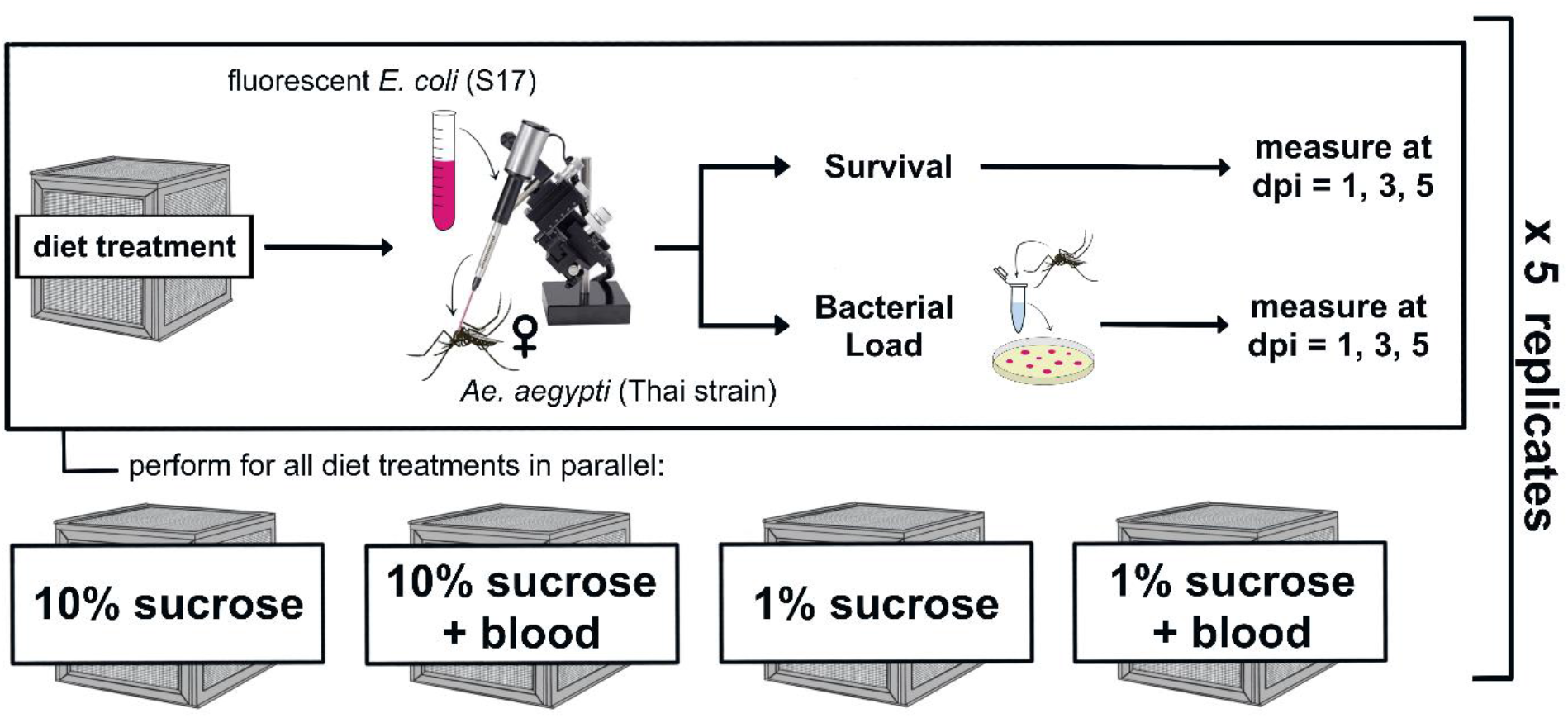
Experimental Design Schematic. We reared adult female *Ae. aegypti* on four experimental diets: 10% sucrose alone, 10% sucrose + blood meal, 1% sucrose alone, and 1% sucrose + blood meal. We then infected individuals with fluorescent *E. coli* (S17) via intrathoracic microinjection. Infected mosquitoes were split into monitoring groups for survival and bacterial load. Survival and bacterial load measurements were obtained in parallel at 1, 3, and 5 days post-infection (dpi). The resultant dataset was used to build models describing the effect(s) of blood and/or sucrose diets on resistance and tolerance simultaneously (Table 1).

### Dietary sucrose and blood affect resistance

#### Lower dietary sucrose increases resistance

We measured resistance to bacterial infection by testing the effects of Sucrose and Blood on Bacterial Load at 1, 3, and 5 days post-infection (dpi). We first built a model using Sucrose, Blood, and Day as predictor variables and Bacterial Load as the response variable, then assessed main effects as well as all potential interactions. We performed backward elimination to identify significant model terms and found that both Sucrose (Table 2: Model A, p _Sucrose_ = 4.57 × 10^−5^) and Day (Table 2: Model A, p _Day_ = 2.92 × 10^−4^) significantly affect resistance to infection. Females fed 1% sucrose had significantly lower bacterial loads (and therefore higher resistance) than females fed 10% sucrose, and bacterial load decreased over time for both treatment groups (Fig 2). Furthermore, pairwise comparisons between sucrose treatments on each individual day showed that 1% sucrose exposure significantly enhanced resistance to infection at all days post-infection. (Table 3: Model B (Day 1), p _Sucrose_ = 0.008; Model C (Day 3), p _Sucrose_ = 0.005; Model D (Day 5), p _Sucrose_ = 0.048).

**Table 2.** Across-Days Resistance Model A and Summary Output.

| Predictors | Bacterial Load |  |  |  |
| --- | --- | --- | --- | --- |
|  | Estimate | SE | Statistic | p-value |
| (Intercept) | 4.01 | 0.98 | 4.10 | 1.42e-4 |
| Day | -1.04 | 0.27 | -3.88 | **2.92e-4** |
| Sucrose | 3.81 | 0.86 | 4.44 | **4.57e-5** |
| Observations | 56 |  |  |  |

Resistance Model A is a linear model testing the effect(s) of Blood, Sucrose, and Day on Bacterial Load.

**Table 3.** Day-Specific Resistance Models B-D and Summary Outputs.

Summary Outputs:
| Resistance Model B (Day 1) |  |  |  |  | Resistance Model C (Day 3) |  |  |  | Resistance Model D (Day 5) |  |  |  |
| --- | --- | --- | --- | --- | --- | --- | --- | --- | --- | --- | --- | --- |
| Bacterial Load |  |  |  |  | Bacterial Load |  |  |  | Bacterial Load |  |  |  |
| *Predictors* | *Est.* | *SE* | *Stat.* | *p-value* | *Est.* | *SE* | *Stat.* | *p-value* | *Est.* | *SE* | *Stat.* | *p-value* |
| (Intercept) | 4.40 | 1.41 | 3.11 | 0.006 | 0.22 | 1.04 | 0.21 | 0.836 | 0.15 | 0.45 | 0.34 | 0.742 |
| Sucrose | 4.86 | 1.63 | 2.98 | **0.008** | 4.70 | 1.47 | 3.19 | **0.005** | 1.39 | 0.64 | 2.17 | **0.048** |
| Blood | -3.67 | 1.63 | -2.25 | **0.038** |  |  |  |  |  |  |  |  |
| Observations | 20 |  |  |  | 20 |  |  |  | 16 |  |  |  |

Resistance Models B-D are linear models testing the effect(s) of Blood and Sucrose on Bacterial Load at each dpi.

**Fig 2.**
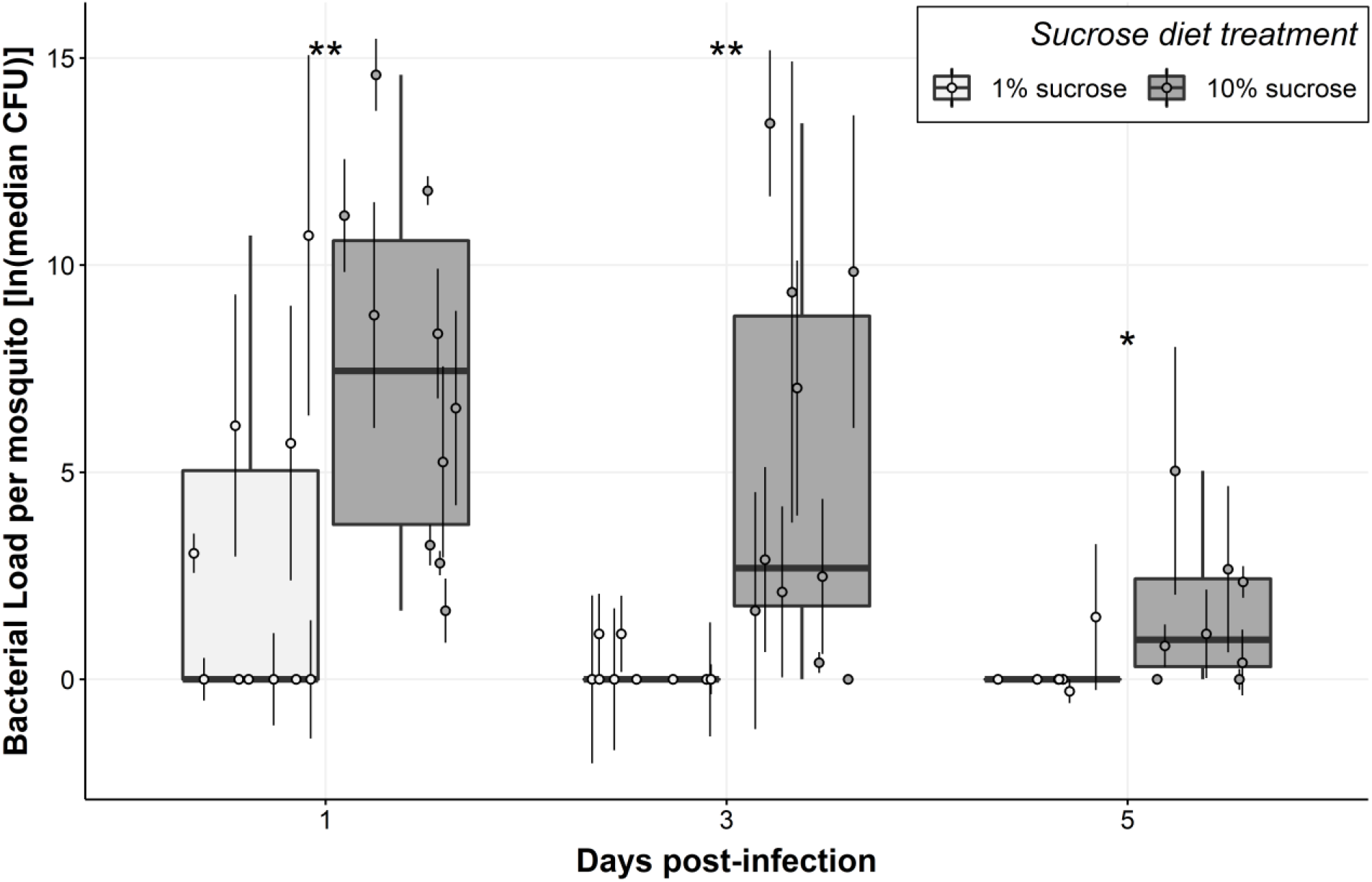
Mosquitoes fed 1% sucrose had significantly lower bacterial loads compared to mosquitoes fed 10% sucrose. Boxplots were constructed using median bacterial loads for females fed 1% sucrose and 10% sucrose at 1, 3, and 5 dpi. Each median value was calculated from four individuals, and point error bars show the interquartile range. Asterisks represent significant differences in bacterial load between sucrose treatments at each time post-infection (** p < 0.01, * p < 0.05). Data were collected from a total of five replicate experiments.

#### Blood increases resistance at 1 dpi only

Blood feeding also significantly affected resistance to infection, but only at 1 dpi, at which blood-fed females had significantly lower bacterial loads compared to non-blood-fed females (Table 3: Model B, p _Blood_ = 0.038; Fig 3).

**Fig 3.**
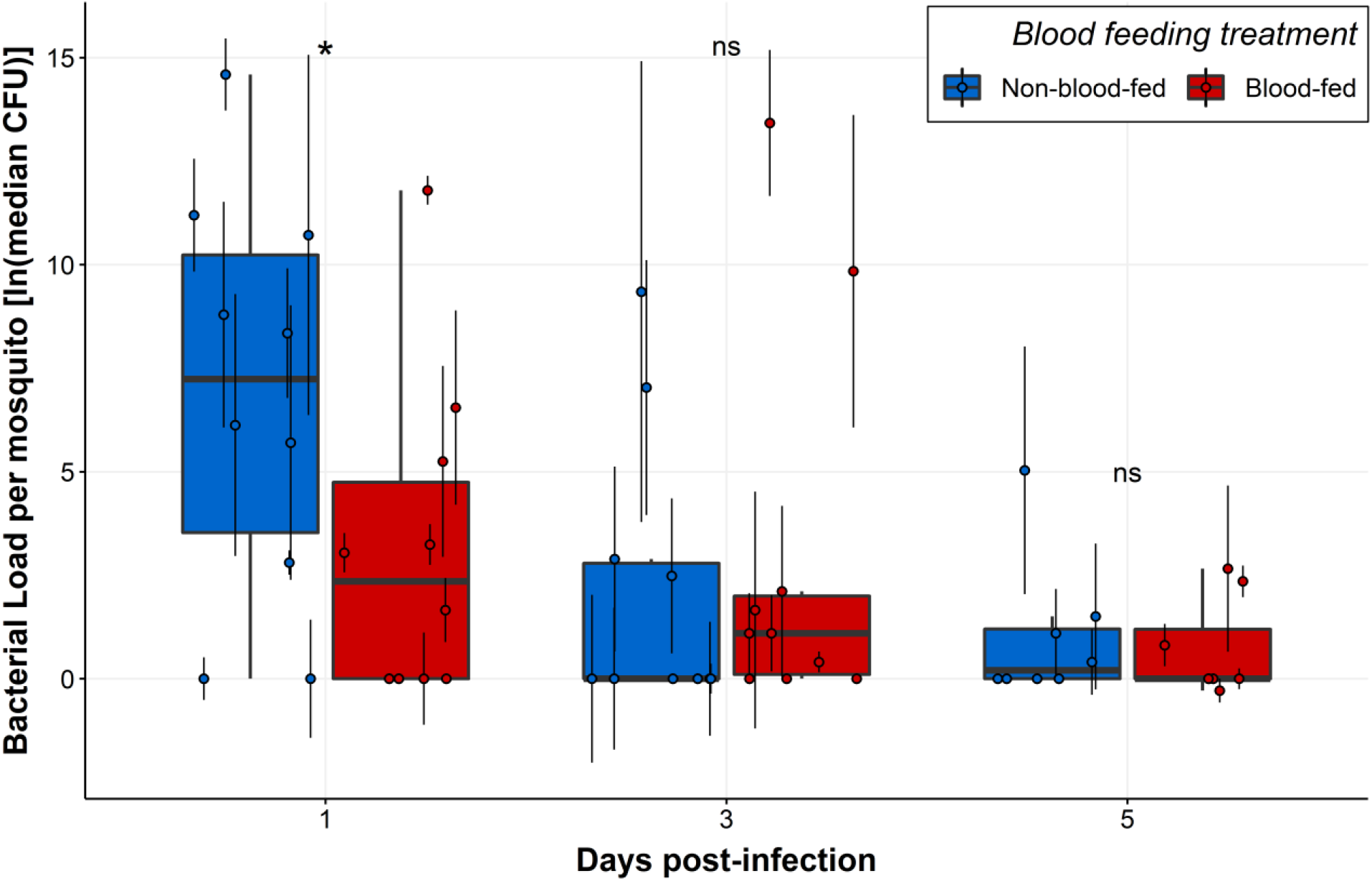
Blood-fed mosquitoes had significantly lower bacterial loads compared to non-blood-fed mosquitoes at 1 dpi. Boxplots were constructed using plotted points showing median (n = 4) bacterial load for blood-fed and non-blood-fed females at 1, 3, and 5 dpi. Each median value was calculated from four individuals, and point error bars show the interquartile range. Asterisks represent significant differences in bacterial load between sucrose treatments at each time post-infection (* p < 0.05, ns = no significant difference). Data were collected from a total of five replicate experiments.

### The effect of a blood meal on tolerance is dynamic across infection timecourse

We also tested the effects of Blood and Sucrose on tolerance to infection, where tolerance was measured as the slope of population survival regressed on pathogen load (24). We first built a model with Survival as the response variable and Bacterial Load, Blood, Sucrose, and Day as the predictor variables. We assessed main effects as well as all potential interactions and performed backward elimination to achieve the best fit model. The final model revealed a significant three-way interaction between Blood, Bacterial Load, and Day (Table 4: Model E; p _Bacterial Load × Blood × Day_ = 0.006), indicating that the effect of blood on tolerance changes across time. In order to examine the nature of blood’s time-dependent effect on tolerance, we parsed our data by day and used the resulting three datasets to build a tolerance model for each day (Fig 4). These day-specific models reveal that blood-fed mosquitoes had significantly lower tolerance at 1 dpi (Table 5: Model F, p _Bacterial Load × Blood_ = 0.002), no significant difference at 3 dpi, and significantly higher tolerance at 5 dpi (Table 5: Model G, p _Bacterial Load × Blood_ = 0.040) when compared to non-blood-fed mosquitoes. In parallel, we also tested whether tolerance changes across time for blood-fed and/or non-blood-fed treatment groups. To achieve this, we parsed our dataset by blood feeding status and used the resulting two datasets to build a tolerance model for each blood feeding treatment group. These models revealed that tolerance significantly decreased over time in non-blood-fed mosquitoes (Table 6: Model H, p _Bacterial Load × Day_ = 6.70 × 10^−6^) but did not significantly change in blood-fed mosquitoes (Table 6: Model I).

**Table 4.** Across-Days Tolerance Model E and Summary Output.

| Tolerance Model E<br>(Across Days): | | $E(\text{Alive}, \text{Dead}) = \varphi[\beta_0 + \beta_1(\text{Day}) + \beta_2(\text{Blood}) + \beta_3(\text{Bacterial Load}) \\ + \beta_4(\text{Day} \times \text{Blood}) + \beta_5(\text{Day} \times \text{Bacterial Load}) \\ + \beta_6(\text{Blood} \times \text{Bacterial Load}) \\ + \beta_7(\text{Day} \times \text{Blood} \times \text{Bacterial Load})]$ | | |
| --- | --- | --- | --- | --- |
| <u>Summary Output:</u> |  |  |  |  |
|  |  | Survival (Alive, Dead) |  |  |
| *Predictors* | *Estimate* | *SE* | *Statistic* | *p-value* |
| (Intercept) | 1.34 | 0.33 | 4.08 | 1.67e-4 |
| Day | -0.17 | 0.08 | -2.05 | **0.046** |
| Blood | 0.93 | 0.49 | 1.91 | 0.062 |
| Bacterial Load | 0.14 | 0.06 | 2.37 | **0.022** |
| Blood : Bacterial Load | -0.24 | 0.09 | -2.59 | **0.013** |
| Day : Bacterial Load | -0.07 | 0.02 | -3.42 | **0.001** |
| Blood : Day | -0.22 | 0.12 | -1.79 | 0.080 |
| Day : Blood : Bacterial Load | 0.09 | 0.03 | 2.86 | **0.006** |
| Observations | 56 |  |  |  |

Tolerance Model E is a quasibinomial generalized linear model testing the effect(s) of Blood, Sucrose, Day, and Bacterial Load on Survival (Alive, Dead).

**Table 5.** Day-Specific Tolerance Models F-G and Summary Outputs.

|  |  |
| --- | --- |
| Tolerance Model F (Day 1): | $P(\text{Alive}, \text{Dead}) = \varphi[\beta_0 + \beta_1(\text{Blood}) + \beta_2(\text{Bacterial Load}) + \beta_3(\text{Bacterial Load} \times \text{Blood})]$ |
| Tolerance Model G (Day 5): | $E(\text{Alive}, \text{Dead}) = \varphi[\beta_0 + \beta_1(\text{Blood}) + \beta_2(\text{Bacterial Load}) + \beta_3(\text{Bacterial Load} \times \text{Blood})]$ |

Summary Outputs:
| Tolerance Model F (Day 1) |  |  |  |  | Tolerance Model G (Day 5) |  |  |  |
| --- | --- | --- | --- | --- | --- | --- | --- | --- |
| *Predictors* | Survival (Alive, Dead) |  |  |  | Survival (Alive, Dead) |  |  |  |
|  | *Estimate* | *SE* | *Statistic* | *p-value* | *Estimate* | *SE* | *Statistic* | *p-value* |
| (Intercept) | 0.93 | 0.23 | 4.10 | 4.19e-5 | 0.48 | 0.20 | 2.36 | 0.036 |
| Blood | 1.20 | 0.39 | 3.10 | **0.002** | -0.21 | 0.30 | -0.71 | 0.492 |
| Bacterial Load | 0.08 | 0.03 | 2.36 | **0.019** | -0.33 | 0.13 | -2.53 | **0.026** |
| Blood : Bacterial Load | -0.17 | 0.06 | -3.08 | **0.002** | 0.52 | 0.23 | 2.31 | **0.040** |
| Observations | 20 |  |  |  | 16 |  |  |  |

Tolerance Model F is a binomial generalized linear model testing the effect(s) of Blood, Sucrose, and Bacterial Load on Survival (Alive, Dead) at 1 dpi. Tolerance Model G is a quasibinomial generalized linear model testing the effect(s) of Blood, Sucrose, and Bacterial Load on Survival (Alive, Dead) at 5 dpi.

**Table 6.** Blood Feeding Status-Specific Tolerance Models H-I and Summary Outputs.

|  |  |  |  |  |  |  |  |  |
| --- | --- | --- | --- | --- | --- | --- | --- | --- |
| Tolerance Model H (Non-Blood-Fed): | $P(\textit{Alive}, \textit{Dead}) = \varphi[\beta_0 + \beta_1(\textit{Day}) + \beta_2(\textit{Bacterial Load}) + \beta_3(\textit{Bacterial Load} \times \textit{Day})]$ | | | | | | | |
| Tolerance Model I (Blood-Fed): | $E(\textit{Alive}, \textit{Dead}) = \varphi[\beta_0 + \beta_1(\textit{Day}) + \beta_2(\textit{Bacterial Load})]$ | | | | | | | |
| <u>Summary Outputs:</u> |  |  |  |  |  |  |  |  |
| Tolerance Model H (Non-blood-fed) |  |  |  |  | Tolerance Model I (Blood-fed) |  |  |  |
| Survival (Alive, Dead) |  |  |  |  | Survival (Alive, Dead) |  |  |  |
| *Predictors* | *Estimate* | *SE* | *Statistic* | *p-value* | *Estimate* | *SE* | *Statistic* | *p-value* |
| (Intercept) | 1.34 | 0.25 | 5.38 | 7.38e-8 | 2.13 | 0.35 | 6.11 | 2.18e-6 |
| Day | -0.17 | 0.06 | -2.70 | **0.007** | -0.35 | 0.08 | -4.17 | **3.23e-4** |
| Bacterial Load | 0.14 | 0.04 | 3.12 | **0.002** | -0.04 | 0.03 | -1.46 | 0.158 |
| Day : Bacterial Load | -0.07 | 0.02 | -4.50 | **6.70e-6** |  |  |  |  |
| Observations | 28 |  |  |  | 28 |  |  |  |

Tolerance Model H is a binomial generalized linear model testing the effect(s) of Sucrose, Day, and Bacterial Load on survival (Alive, Dead) in non-blood-fed mosquitoes. Tolerance Model I is a quasibinomial generalized linear model testing the effect(s) of Sucrose, Day, and Bacterial Load on survival (Alive, Dead) in blood-fed mosquitoes.

**Fig 4.**
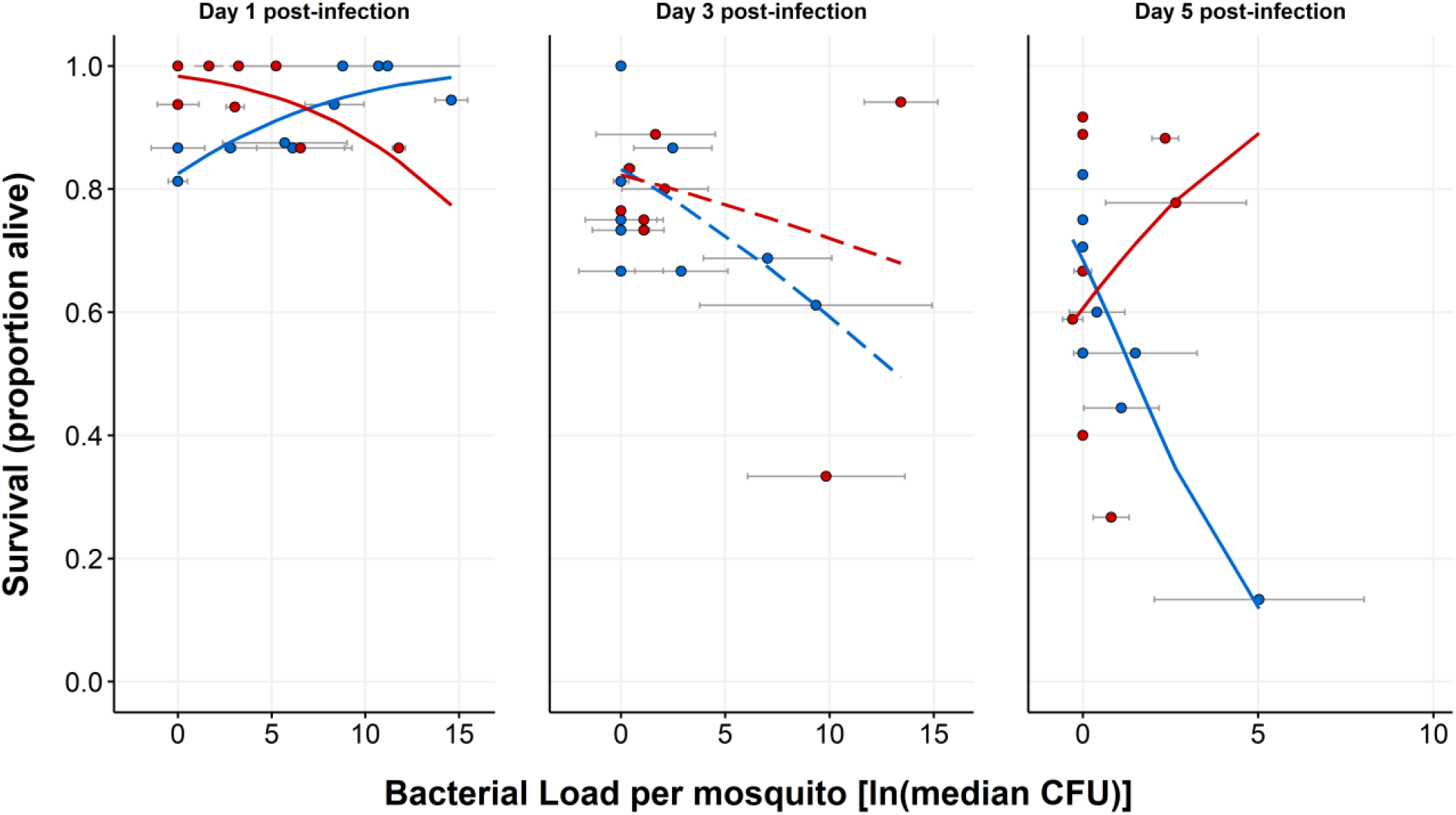
The effect of blood feeding on tolerance varies across time. Points show population survival plotted against median bacterial load at each timepoint post-infection. Error bars around points show the interquartile range of the four values used to calculate bacterial load median values. Plotted lines are derived from interaction plots of estimated marginal means calculated from day-specific binomial tolerance models (Table 5: Model F (Day 1), p _Bacterial Load × Blood_ = 0.002; Day 3, no significant predictors; Table 5: Model G (Day 5), p _Bacterial Load × Blood_ = 0.040) plotted on a response scale rather than a linear scale. The presence of solid lines indicates significantly different slopes, and therefore a significant difference in tolerance, between blood treatment groups at that timepoint. Dashed lines indicate no significant difference in tolerance at that timepoint. Data were collected from a total of five replicate experiments.

### A blood meal alters the shape of a mosquito’s immune defense trajectory through time

Blood altered resistance early in infection and tolerance dynamically across the infection timecourse. Therefore, we compared the relative utilization of each component of immune defense by quantitatively plotting resistance and tolerance for each blood feeding status group at all three timepoints (Fig 5). Blood-fed mosquitoes displayed an increase in resistance (Table 2: Model A, p _Day_ = 2.92 × 10^−4^) and a non-significant trend toward increased tolerance (Table 6: Model I) from 1 dpi to 5 dpi, while non-blood-fed mosquitoes displayed increased resistance (Table 2: Model A, p _Day_ = 2.92 × 10^−4^) but decreased tolerance (Table 6: Model H, p _Bacterial Load × Day_ = 6.70 × 10^−6^) from 1 dpi to 5 dpi (Fig 5).

**Fig 5.**
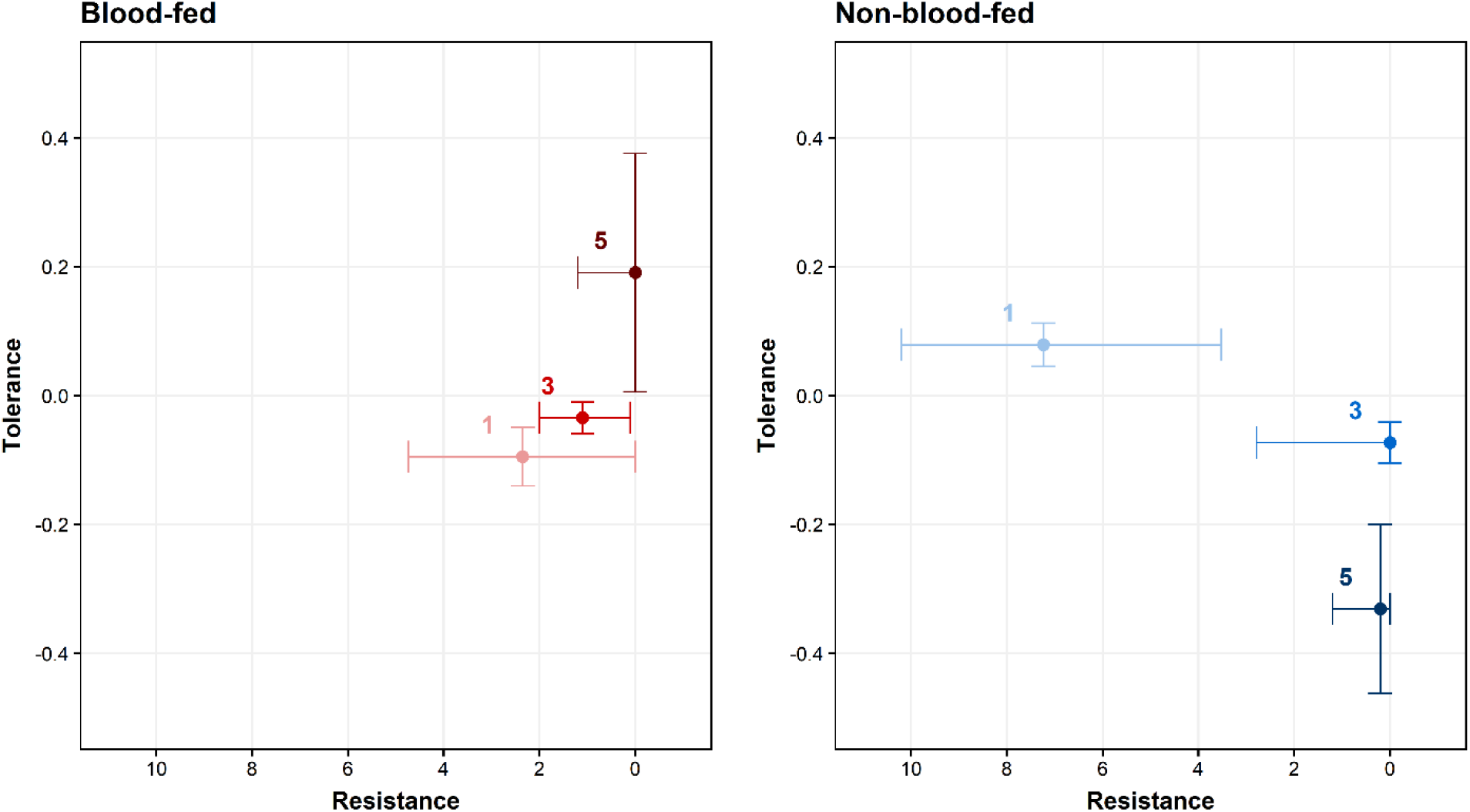
Blood feeding alters the mosquito’s trajectory of resistance and tolerance over time. Plotted points display the balance of tolerance and resistance through an infection timecourse for blood-fed (left panel) and non-blood-fed mosquitoes (right panel). To estimate tolerance, we regressed Bacterial load on Survival separately for each blood feeding status/day combination and found the estimated marginal mean and standard error of Bacterial Load for each model using the emmeans package (25). Resistance is the population median CFU with interquartile ranges for each group plotted on a reversed x-axis scale (as resistance is inversely related to CFU). Data were collected from a total of five replicate experiments.

## Discussion

### Sucrose restriction enhances resistance throughout infection timecourse

We have shown that a 1% sucrose diet confers higher resistance to intrathoracic infection by the gram-negative bacterium *E. coli* S17 in adult female *Ae. aegypti* (Thai strain) compared to a 10% sucrose diet. In mosquitoes, effects of dietary sugars on infection response have primarily been studied using oral infection models. In one study, pre-infection sugar feeding enhanced *Ae. aegypti* resistance to Zika by increasing the expression of antiviral genes (26), while in *An. stephensi*, lower dietary glucose abundance correlated with higher resistance to *Plasmodium berghei* (21). In *D. melanogaster*, lower dietary sugar concentration increased resistance to bacterial infection in two separate studies (22,27). Altogether, the current body of work suggests lower dietary sugar levels usually correlate with increased resistance to infection, which is consistent with our findings.

While dietary sucrose significantly affected mosquito resistance in our study, it did not affect survival (S1 Fig). Conversely, effects of dietary sugar on survival have been observed in *D. melanogaster*, although the directionality of the effect is not consistent. Howick & Lazzaro (22) showed that higher dietary sugars were associated with lower survival (and lower resistance) post-infection. However, other studies have shown a positive relationship between dietary sugar and infection outcome (although resistance was not directly measured). For example, in one study, flies that consumed a relatively low-protein, high-carbohydrate diet displayed increased survival to bacterial infection and higher constitutive expression of antimicrobial peptides (AMPs) (28). In another study, dietary glucose supplementation led to significantly improved *D. melanogaster* longevity and survival following bacterial infection (29). Multiple dissimilarities between these studies and our own, including experimental design and pathogen type, could underlie the different findings. Overall, relationships between dietary sugars and survival after infection are not fully consistent and warrant further study.

The mechanisms by which sugar affects resistance are not known. As diet is known to affect hemolymph composition in insects (30,31), higher dietary sugar may provide a more suitable environment for bacteria to proliferate. Alternatively, but not mutually exclusively, a lower concentration of dietary sucrose may directly or indirectly affect signaling of the mosquito’s immune pathways, thereby inhibiting resistance mechanisms such as melanization, AMP production, and/or hemocyte activity. Higher dietary sugar levels are associated with a stronger melanization response in mosquitoes (32–34), which may initially appear antithetical to our results. However, *E. coli* is preferentially phagocytized rather than melanized by mosquitoes (35–37), suggesting that any reduction in melanization that may have occurred in the 1% sucrose treatment is unlikely to impact infection outcomes in our system. Sucrose may have a different effect after challenge with a pathogen that is primarily melanized. In *D. melanogaster*, differences in dietary sugar levels are associated with physiological changes that may explain effects on resistance. For example, flies reared on high sugar diets display abnormal hemocyte morphology, defects in phagocytosis ability, and excessive activation of Toll/JNK in the fat body (38). Other work has also implicated high-sugar diets in inhibition of FOXO signaling in *D. melanogaster* (39), which has been shown to be critical in resistance to bacterial infection (40).

While the differences in immune resistance between our sucrose treatments may be explained by the effect of dietary sucrose availability *per se*, it is also feasible that the effect we observe is the result of general caloric restriction. Insects subject to food restriction and insects subject to infection undergo many of the same physiological changes, including reconfiguration of intermediate metabolism, reduced energy storage, release of glucose and fatty acids from existing energy stores, and inhibition of the insulin-like signaling pathway (41,42). By virtue of this commonality, food restriction prior to infection may induce a metabolic switch that functionally primes the mosquito to resist pathogen invasion. Indeed, positive effects of starvation or dietary restriction on resistance and/or infection outcome more generally have been observed in multiple insect orders (43–48). Imposed starvation does not always lead to increased resistance, however, and in many cases has been shown to cause decreased resistance and/or adverse infection outcomes (45,49–53). For example, starved *Leptinotarsa decemlineata* beetles display increased susceptibility to *Beauveria bassiana* and heightened mortality post-infection compared to their non-starved counterparts (50). Similarly, the tsetse fly *Glossina morsitans morsitans* displays increased susceptibility to both *Trypanosoma congolense* or *Trypanosoma brucei brucei* under starvation conditions (51).

### Blood improves resistance early post-infection

Our data also reveal that a blood meal enhances resistance at 1 dpi. This differs from the effect of sucrose, which remains significant at every timepoint measured. Previous work reported similar effects of blood feeding on resistance to *E. coli*, in association with insulin signaling in one instance (23) and 20-hydroxyecdysone (20E) in another (54). In addition to supporting these findings, our data contribute an enhanced understanding of the dynamic nature of blood’s effect over time. Specifically, the effect of blood on resistance is relatively transient, especially in comparison to the effect of dietary sucrose on bacterial resistance which is consistent for at least five days post-infection. As the effect of blood on 20E titers generally does not extend past 48 hours post-blood meal (55,56) and 20E can enhance resistance in mosquitoes (Reynolds et al., 2020; but see Wang et al., 2022; Werling et al., 2019), it is possible that the dynamic effect of blood feeding on resistance we observed is due to changing 20E titers over time. Titers of another hormone, Juvenile Hormone (JH), rapidly and drastically decline in blood-fed *Ae. aegypti*, reaching minimums between 24 and 48 hours post-blood meal (59,60). JH and JH analogs are associated with various immunosuppressive effects, including the downregulation of AMP genes (61–63), decreased activity of hemocyte activator molecules (64), and reductions in phenoloxidase activity (65,66).

Therefore, it is also feasible that improved resistance after blood feeding is mediated via declines in JH. In addition to the potential regulatory role of hormones, investigations into the mosquito’s transcriptional profile post-blood feeding indicate altered expression of immune-related genes within the 24 hours following a blood meal (67–70). Such genes may affect resistance to bacterial infection and may or may not be subject to regulation by insulin signaling and/or hormones such as JH and 20E.

### A blood meal, but not sucrose concentration, affects tolerance to infection

While the effect of blood on resistance is limited to 1 dpi, we observe a dynamic effect of blood feeding on tolerance across the 5-day infection timecourse. Blood-fed mosquitoes have significantly lower tolerance compared to non-blood-fed mosquitoes at 1 dpi, and significantly higher tolerance at 5 dpi. Further, non-blood-fed mosquitoes show a significant decrease in tolerance across a 5-day infection timecourse while blood-fed mosquitoes show static tolerance across a 5-day infection timecourse. This suggests that the blood meal ameliorates a decline in tolerance that mosquitoes experience when provided only sugar meals. Blood is a unique nutritional resource – its digestion is associated with various physiological stressors, including rapid shifts in temperature and pH (15,71) as well as heme toxicity and oxidative stress (72–74). Despite the challenges posed by blood digestion, mosquitoes are well adapted to overcome the associated stressors (15,17,71,75,76). Because many stressors associated with blood feeding are also associated with infection (20,21,77,78) it is possible that the homeostasis-promoting processes that occur during or after blood feeding have the additional effect of promoting host health during infection (i.e. tolerance).

Blood feeding induces multiple signaling cascades that may promote tolerance to infection. For example, heat shock proteins (HSP) are widely conserved in both eukaryotes and prokaryotes (79) and are implicated in a variety of processes generally related to the reduction of stress (80). They are rapidly induced upon blood feeding in multiple arthropods and are critical to surviving the stress of a blood meal (15,19,81). Host HSPs have also been implicated in host response to bacterial, viral, and fungal infections alike, playing roles in homeostasis via maintenance of protein stability and functionality, reductions in inflammation, and attenuation of autoimmunity (82,83). HSPs that protect arthropods from blood-induced damage may also promote host health and survival during infection. Additionally, the unfolded protein response (UPR) is a similarly highly conserved, stress-mitigating, pathway that has been heavily implicated in defense against pathogens (reviewed by Rosche et al., 2021). Transient UPR upregulation occurs after blood feeding in *Ae. aegypti* (85). Further, UPR upregulation promotes tolerance to *Pseudomonas aeruginosa* in the nematode *Caenorhabditis elegans* (86). It also promotes survival in mosquito cells infected with dengue virus in vitro by ameliorating infection-induced endoplasmic reticulum stress (87), a function which is characteristic of tolerance. Importantly, it is unlikely that any single signaling pathway or molecular cascade regulates tolerance to infection, as immunopathology affects various physiological processes (1) that require repair or protection. Thus, if the aforementioned processes do indeed affect tolerance, they likely do so in tandem with other mechanisms.

It is also possible that the tolerance benefit of a blood meal is explained by the influx of nutrients (e.g., protein, lipid) associated with a blood meal. However, we observed no effect of sucrose on tolerance, and higher dietary sucrose concentrations are also strongly associated with higher nutrient reserves. Whole-body homogenates of mosquitoes fed 10% sucrose have significantly higher levels of sugar, glycogen, and lipids compared to mosquitoes fed 2% sucrose (88). This indicates that if tolerance is indeed regulated nutritionally in our system, this regulation may be specific to the nutrients provided by a blood meal.

### Blood-fed and non-blood-fed mosquitoes employ different, but equally effective, immune defense strategies

Blood-fed and non-blood-fed mosquitoes showed markedly different resistance-tolerance strategies for surviving infection with a non-coevolved bacterial pathogen (Figure 5). In the absence of a blood meal, a mosquito displays an increase in resistance and a decrease in tolerance across a 5-day infection timecourse. But when provided with a blood meal, early infection is characterized by a significantly stronger resistance response and significantly weaker tolerance response (possibly the result of a trade-off). Across time, blood-fed mosquitoes show no change in tolerance, and an increase in resistance that is similar to that of non-blood-fed mosquitoes. In comparing the groups at the end of infection timecourse, blood-fed mosquitoes show equal resistance and higher tolerance compared to non-blood-fed mosquitoes. Each diet clearly induced a unique immune defense strategy. Interestingly, we observed no significant difference in survival (S1 Fig) indicating that the two strategies employed by the groups are equally effective in surviving infection. A blood meal is not only physiologically stressful and energetically costly to digest on its own (89), it also catalyzes reproductive processes (i.e. vitellogenesis) that are energetically costly (90). Immune defense is also a costly process (10,91–93). Understanding how organisms partition limited resources amongst energy-intensive processes represents an ongoing challenge in biology. Reproduction, immunity, and digestion have been shown to physiologically trade off in multiple insect systems (90,94–100). In light of this, it may be somewhat surprising that mosquitoes under the combined stress of blood digestion, reproduction, and infection in parallel show no difference in survival compared to non-blood-fed mosquitoes. Overall, our results therefore suggest that the immune defense benefits of a blood meal are great enough to mitigate the stressors and resource usage associated with this meal.

## Conclusions

In the present study, we show the effects of blood ingestion and two dietary sucrose concentrations on both resistance and tolerance to the non-coevolved bacterium *E. coli* in the adult female yellow fever mosquito *Ae. aegypti*. Our results indicate that dietary sucrose concentration and blood ingestion both affect resistance, while only blood ingestion affects tolerance. The effect of blood on tolerance was dynamic across time, significantly worsening tolerance at the start of the infection timecourse and significantly enhancing tolerance by the end. Mosquitoes are one of many arthropods that transmit pathogens by virtue of a hematophagous lifestyle: mosquitoes, ticks, biting midges, triatomine bugs, fleas, black flies, and sand flies alike all transmit pathogens when consuming vertebrate blood (101). Likewise, they all share the ability to tolerate the pathogens they transmit, which is critical for successful transmission. In light of this, motivation for exploring the relationship between hematophagy and tolerance is not limited to mosquitoes, but rather, is broadly relevant in all arthropod-borne pathogen systems. Further, shared characteristics of tolerance biology could potentially be leveraged as novel targets for effective and sustainable vector control (102). Because blood feeding is already one of the best-described areas of mosquito biology, has been investigated (103–106) and implemented (107) as a vector control target, and affects tolerance in *Ae. aegypti*, targeting blood feeding in the context of tolerance-focused vector control holds excellent potential.

## Methods

### Mosquitoes

Throughout the duration of the experiment, *Ae. aegypti* Thai strain (Laura C. Harrington, Cornell University) mosquitoes were reared in a chamber maintained at 27°C and 80% relative humidity under a 14h:10h light:dark cycle. First, eggs were hatched in RO water placed in a vacuum chamber. Upon hatching, larvae were reared in trays containing RO water at a density of 200-300 larvae per tray and given one pinch of Tetramin fish flakes as well as cat food ad libitum until pupation. Upon pupation, pupal cups were split into four 8”x8” mesh treatment cages (Bioquip, Rancho Dominguez, CA, USA): 10% sucrose only, 10% sucrose + blood, 1% sucrose only, and 1% sucrose + blood. Each cage received the appropriate sucrose meal concurrent with the addition of pupal cups. Pupae were allotted 48 hours to eclose before pupal cup removal. Post-eclosion, all adults were left undisturbed for 48 hours, then starved for 24 hours. Next, mosquitoes from cages containing blood feeding treatments were provided with a blood meal maintained at 37 °C via a membrane feeding system (Hemotek, Blackburn, UK) for 1-2 hours. Sucrose meals were then returned to all mosquitoes. Infections were performed on females at 24 hours post-blood feeding. At this time, blood feeding status was confirmed visually under a microscope by the presence of a blood bolus.

### Sucrose and blood diets

Mosquitoes were maintained on one of four experimental diets: 10% sucrose alone, 10% sucrose + blood, 1% sucrose alone, and 1% sucrose + blood. Sucrose meals were created by passing a solution of DI water and UltraPure sucrose (Life Technologies, Carlsbad, CA) through a 0.2 μm filter. Blood meals consisted of defibrinated rabbit blood (Hemostat, Dixon, CA, USA) supplemented with Na_2_ATP to a 1 mM concentration.

### Bacterial culture and mosquito infections

The bacteria used for infections, *E. coli* S17 pPROBE-mCherry (Dimopoulos Lab, unpublished data), contains a fluorescent mCherry plasmid and a kanamycin resistance cassette. Bacteria were grown in Luria broth (LB) supplemented with kanamycin (50 μg/mL) overnight at 30°C with shaking. Cultures were washed thrice in sterile 1X PBS, then pelleted and resuspended to OD600 = 1 ± 0.1 (mean of 1.23×^9^ CFU/mL across replicates). At the time of infection, mosquitoes were anesthetized on ice and females were injected with 69 nanoliters of a 1 × 10^−2^ dilution of this culture by piercing the soft tissue of the anepisternal cleft of the mesothorax using a Nanoject II Auto-Nanoliter Injector (Drummond, Broomall, PA, USA). Fresh injection needles were prepared for each replicate by manually pulling a borosilicate glass capillary tube (Drummond, Broomall, PA, USA) over a flame to achieve a tip with an outer diameter no greater than 500 microns as measured using a stage micrometer. Preliminary experiments indicated that Nanoject II delivery of bacteria was adequately precise, delivering a mean of 312 ± 13 CFU per mosquito (n = 6 mosquitoes were injected, then each homogenate was immediately cultured as described in the subsequent section) (S1 File). Following infection, mosquitoes from each diet group were randomly allocated into two groups and placed in separate cages for survival and bacterial load measurements.

### Monitoring survival and bacterial load

Survival and bacterial load were measured on days 1 (24 hours +/-2 hours), 3 (72 hours +/-2 hours), and 5 (120 hours +/-2 hours) post-infection. Immediately after data collection at each timepoint, dead individuals were removed from survival and bacterial load cages and discarded. Survival was monitored by counting the number of mosquitoes dead and alive. To measure bacterial load, four living mosquitoes were sampled using an InsectaVac Aspirator (Bioquip, Rancho Dominguez, CA, USA) and individually homogenized in 150 μL sterile 1X PBS. Serial dilutions were performed at 10^−2^ and 10^−4^ and cultured alongside undiluted homogenate on LB supplemented with kanamycin (50 μg /mL) at 30°C for 24-48 hours. Resulting fluorescent colonies were counted using a stereo microscope fluorescence adapter system (Nightsea, Lexington, MA). For each individual, the least dilute plate with countable CFUs was used to obtain a representative value. After obtaining a CFU count for each individual mosquito, the median of four mosquitoes was calculated and used in the data set. Each row in the data set (S1 File) contains the median CFU of four mosquitoes paired with the accompanying survival value for that treatment group.

### Statistical analysis

All analyses were performed using R statistical software (108) and RStudio (109).

To measure resistance, we built linear models testing the effect(s) of Blood, Sucrose, Day, and Replicate on Bacterial Load (Table 1). Backward elimination was used to assess all possible interactions as well as main effects. The presence of any main effect indicates that variable significantly affects resistance.

To measure tolerance, we used a reaction norm approach (24) to test for variation in health across a range of real-time pathogen loads between diet groups at multiple time points. Our approach is thus categorized as range tolerance rather than point tolerance (110). We built binomial generalized linear models testing the effect(s) of Blood, Sucrose, Day, and Bacterial Load on Survival (Table 1). Backward elimination was used to assess all possible interactions as well as main effects. Further, we tested all tolerance models for overdispersion (111) by performing residual deviance goodness-of-fit tests and Pearson goodness-of-fit tests. When detected, we adjusted for overdispersion by correcting the standard errors in those models through incorporation of a dispersion parameter when calculating the variance. Models with overdispersion parameters are defined as quasi-generalized linear models using a binomial distribution where the variance is given by φ × μ, in which μ is the mean and φ the dispersion parameter. We used these final models to interpret any effect(s) of our predictor variables on tolerance. Specifically, a significant interaction between Bacterial Load and another variable in predicting Survival indicates that variable significantly affects tolerance.

To obtain interaction plots of estimated marginal means for each day-specific binomial tolerance model (Figure 4), we used the emmip function from the emmeans package (25). To obtain representative values for tolerance for each level of Blood on Days 1, 3, and 5 (Figure 5), we used the emtrends function from the emmeans package, a function that creates a reference grid for each model of interest, then calculates difference quotients of predictions from those reference grids and computes the marginal averages and standard errors for those averages. We used the emtrends function to obtain these values for each day using Model F (Day 1), Model G (Day 5), and a model with a non-significant Bacterial Load × Blood term for Day 3 (model not shown).

## Acknowledgements

We thank Ellen Klinger and Megan Meuti (The Ohio State University) for statistical advice and helpful experimental design comments, George Dimopoulos and Yuemei Dong (Johns Hopkins University) for providing the mCherry-transformed strain of *E. coli*, as well as Laura C. Harrington (Cornell University) for the gift of *Ae. aegypti* (Thai strain).

## Supporting information

**S1 Fig. Survival across a 5-day infection timecourse is affected by neither blood nor sucrose**. Kaplan-Meier curves are displayed to compare the effects of (A) blood treatment and (B) sucrose treatment on survival in mosquitoes infected with *E. coli*. Corresponding log-rank test p-values are displayed on each panel.

**S1 File. Selected R code used for statistical analysis and figures**.

**S1 Dataset. Data used in analyses**. Data are divided into three tabs: final dataset (df_final), survival analysis dataset (df_surv), and raw CFU dilution counts (raw CFU counts) used to calculate medians used in df_final.

