## Supplementary figures and images for "Sugar restriction and blood ingestion shape divergent immune defense trajectories in the mosquito *Aedes aegypti*"

### S1 Fig

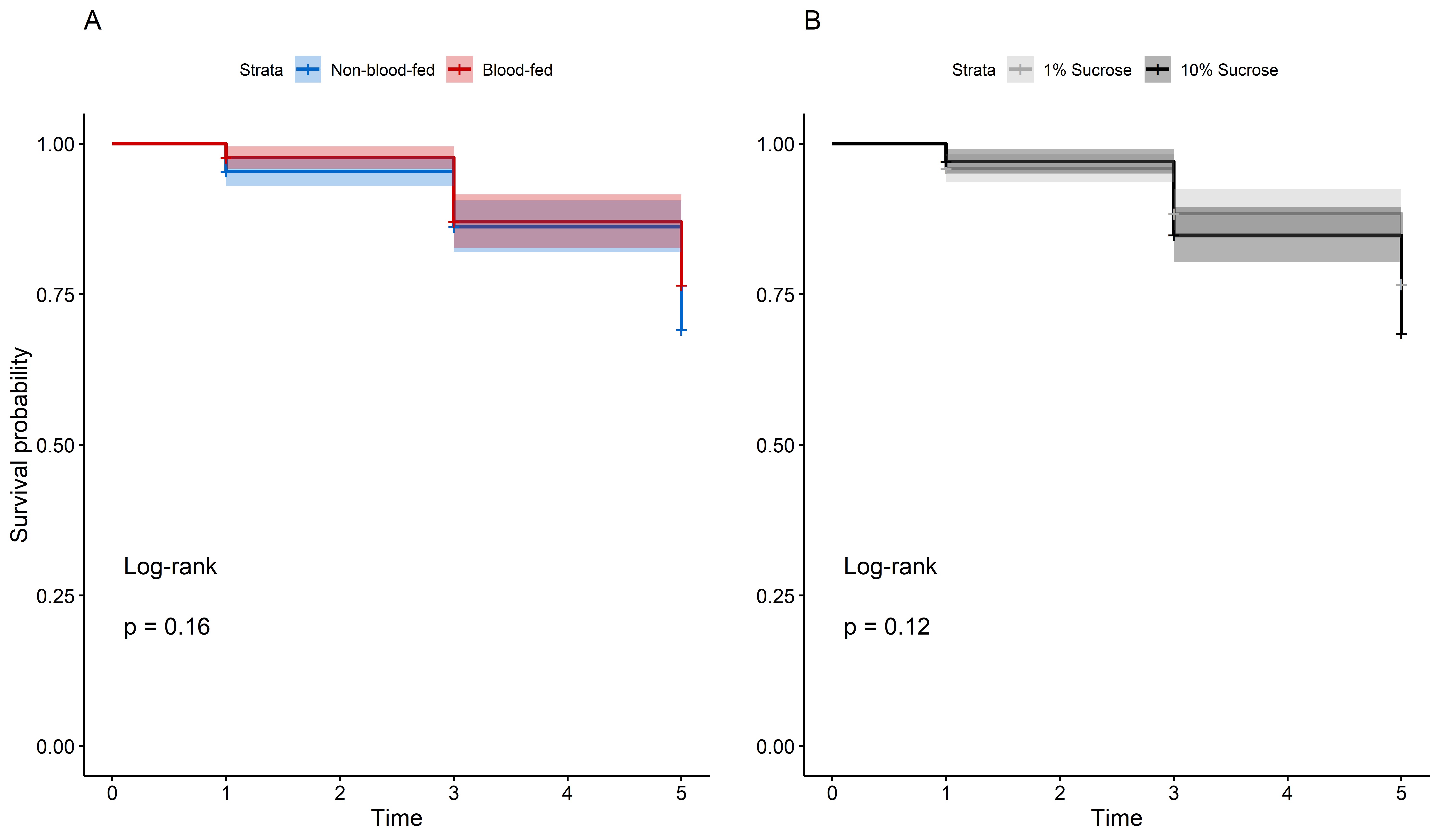
